# Computational investigation of protein photoinactivation by molecular hyperthermia

**DOI:** 10.1101/2020.07.22.216069

**Authors:** Peiyuan Kang, Chen Xie, Oumar Fall, Jaona Randrianalisoa, Zhenpeng Qin

**Author notes:** These authors contributed equally to this work (Peiyuan Kang, Chen Xie).

## Abstract

To precisely control protein activity in a living system is a challenging yet long-pursued objective in biomedical sciences. Recently we have developed a new approach named molecular hyperthermia (MH) to photoinactivate protein activity of interest without genetic modification. MH utilizes nanosecond laser pulse to create nanoscale heating around plasmonic nanoparticles to inactivate adjacent protein in live cells. Here we use a numerical model to study important parameters and conditions for MH to efficiently inactivate proteins in nanoscale. To quantify the protein inactivation process, the impact zone is defined as the range where proteins will be inactivated by nanoparticle localized heating. Factors that reduce the MH impact zone include stretching the laser pulse duration, temperature-dependent thermal conductivity (versus constant properties), and non-spherical nanoparticle geometry. In contrast, the impact zone is insensitive to temperature-dependent material density and specific heat, as well as thermal interface resistance based on reported data. The low thermal conductivity of cytoplasm increases the impact zone. Different proteins with various Arrhenius kinetic parameters have significantly different impact zones. This study provides guidelines to design the protein inactivation process in MH.

## Introduction

Using light to selectively and remotely control biological functions of living systems is a long-pursed objective in biomedical science [1]. Various approaches have been developed during the last decade, such as optogenetics[2] and synthetic photoswitches[3, 4]. Optogenetics utilizes photosensitive molecules to genetically modify the protein of interest and enables light manipulation of targeted cells. This revolutionary technique helps us understand how specific cell type contributes to complex neural circuits and brain functions. While optogenetics is undoubtedly powerful, the requirement of genetic modification still limits its clinical and translational value for human. Another emerging approach to control protein activity with light is by synthetic photoswitches. However, there are limited numbers of photoswitches (mostly based on azobenzene) available, thus narrow options of wavelength window. To overcome these drawbacks, nanoparticles provide promising opportunities in this area [5-8]. One attractive candidate is the plasmonic nanoparticle [9]. When metal particle size is in the nanoscale, nanoparticles exhibit unique optical properties known as plasmonic nanoparticles due to the strong coupling between light and electron oscillation. Among different plasmonic nanoparticles, gold nanoparticles, due to the good chemical stability, have been used to optically control tissue and cellular behaviors, including photothermal therapy of tumor treatment[10], neuron firing[11-13], heat shock protein expression[14] and optoporation[15, 16].

Upon light irradiation, gold nanoparticles convert electromagnetic wave energy into thermal energy and heat up surrounding medium, known as plasmonic heating [17, 18]. The plasmonic heating process can be engineered on different time and length scales by controlling the light energy input [19, 20]. This is because the heat diffusion is highly related to time. A long duration of energy input (such as seconds to minutes) allows the heat to dissipate away from individual particles. The collective heating effect can lead to a global temperature rise in large scales and be used in various biomedical applications, such as thermal therapy for cancer (Fig.1) [21]. On the other hand, the localized heating can be achieved by applying an intense energy input with a short duration (such as a nanosecond laser pulse). In this scenario, the short pulse duration locally heat up the nanoparticle and surrounding medium but is insufficient to heat up the entire system. Hence, highly localized heating can be achieved around individual particles. By tuning the laser pulse duration, the localized heating area can be in the nanoscale (Fig.1). The nanoscale plasmonic heating provides new approaches to trigger cellular or even subcellular thermal responses [19, 22-24].

Recently, we developed a method called molecular hyperthermia (MH) to photoinactivate protein activity [25, 26] by taking advantage of nanoscale plasmonic heating. MH utilizes nanosecond laser pulses to excite plasmonic nanoparticle, for example gold nanosphere (AuNS), as a nanoheater to thermally inactivate proteins adjacent to the nanoparticle. MH selectively inactivates enzymes targeted by AuNS and leaves untargeted enzymes intact, demonstrating the spatial selectivity of MH [25, 27]. We further showed that MH can be used to inactivate cell membrane proteins in live cells without compromising cell viability [26]. MH is a promising method to selectively and remotely manipulate protein activity and cellular behavior. However, MH is a complicated process, and involves not only heat transfer but also temperature-dependent chemical reactions (thermal inactivation of proteins). So far, most models of nanoparticle plasmonic heating focus only on heat transfer process [23, 28, 29]. To better understand the MH, a numerical model would allow quantitative description of protein inactivation induced by nanosecond laser plasmonic heating.

In this work, we developed a numerical model that can quantitatively describe the confined nanoscale heating of gold nanoparticle and protein inactivation during MH. With this model, we further investigated the important parameters for MH, such as laser pulse shape, temperature-dependent material properties, and particle shape. This work provides a better understanding of the protein inactivation response to the nanoscale plasmonic heating, and design guidelines for biomedical applications of MH.

## Method

### Heat transfer model and analytical solution

The analytical solution of spherical nanoparticle heat transfer is used as a benchmark for numerical solution. The model is a spherical nanoparticle as a heat source immersed in aqueous medium. The heat generation is homogeneous inside the nanoparticle and two-temperature effect is negligible in this study. The reason is that the electron-electron interaction time (femtoseconds) and electron-phonon thermalization time (picoseconds) are very short compared with laser pulse duration (nanoseconds) [30]. We neglect the interfacial thermal resistance in the analytical solution, and we discuss the effect of the thermal interfacial resistance in numerical solution.

The heat transfer governing equations is given by Equ. 1-2

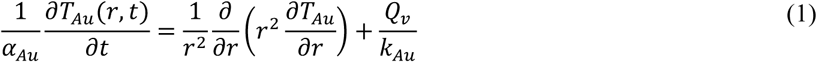

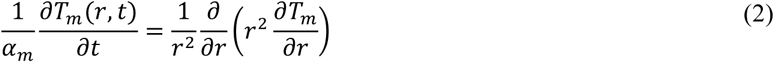

where k is thermal conductivity. α is thermal diffusivity. the subscript “m” and “Au” indicate medium or gold. Upon laser irradiation, the light energy is firstly absorbed by the electrons at surface and the thermal equilibrium of electron gas in the whole nanoparticle will be soon reached due to electron-electron scattering. Next, energy is transferred from electron gas into lattice. Because the electron-electron interaction is much faster (in fs range) than lattice heating (in ps range), the energy transfer to lattice is uniform in the nanoparticle and can be considered as volumetric heating (Q_v_). The rectangular pulse induced volumetric heat source of nanoparticle is expressed as the equation below:

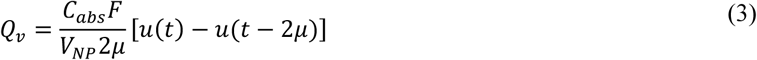

u(t) is unit function, *μ* is central time of pulse (5 ns), and *2μ* is pulse duration (10 ns). F is the laser fluence (25.4×10^9^ W m^-2^). C_abs_ is the absorption cross section of AuNS, calculated by Mie theory (1.38×10^−15^ m^2^)[31]. V_NP_ is volume of AuNS.

The boundary conditions and the initial condition are given be Equ. 4-6.

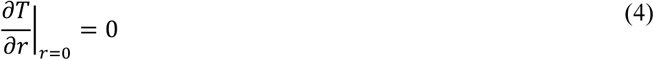

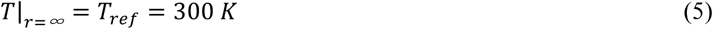

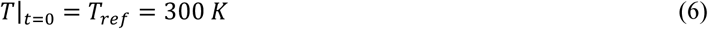

At the interface of gold and medium, we neglect the interfacial thermal resistance, and the continuous interfacial condition is:

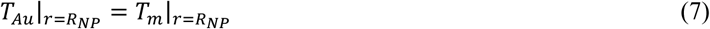

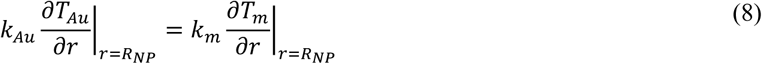

where R_NP_ is radius of nanoparticle (15 nm).

The analytical solution was in part modified from Goldenberg *et al*. and we added the cooling process [32]. First we changed the variable as *V*_1_ *= T*_*Au*_ − *T*_*ref*_ and *V*_*2*_ *= T*_*m*_ − *T*_*ref*_. Second, we introduced Laplace transformation 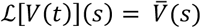 to Equ 1-8:

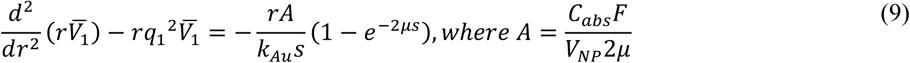

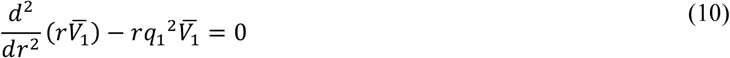

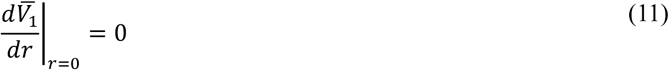

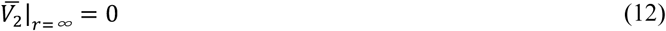

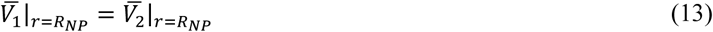

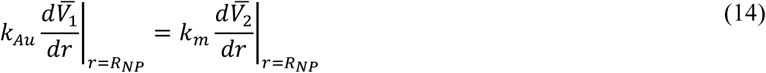

where *q*_1_^*2*^ *= s/α*_*Au*_ and *q*_*2*_^*2*^ *= s/α*_*m*_.

Third, the solution for Equ. (9)-(14) was derived from Goldenberg et al.:

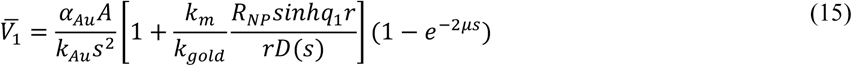

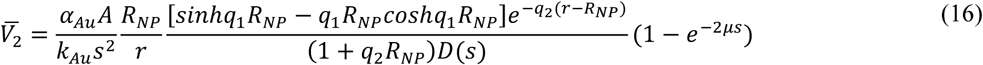

where 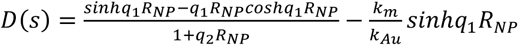.

Next, we inversed Laplace transform the Equ. (15) and (16) with 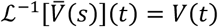. Let 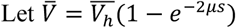. Goldenberg et al. gave the solution of heating process as 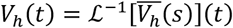 (Equ. 18-20). The whole solution considering both heating and cooling process is given by:

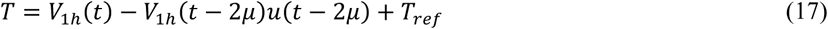

The expansion of analytical solution (Equ. 17) is organized in the table below.

**Table 1.** Analytical solution of single gold nanosphere heated by a rectangular pulse.

|  |  |
| --- | --- |
| Heating process | $T_{Au}(r, t)\vert _{t < 2\mu} = T_{ref} + \frac{R_{NP}^2 C_{abs} F}{k_{Au} V_{NP} 2\mu} \left\{ \frac{1}{3} \frac{k_{Au}}{k_m} + \frac{1}{6} \left( 1 - \frac{r^2}{R_{NP}^2} \right) - \frac{2R_{NP} b}{r\pi} \int_0^\infty \frac{\exp(-y^2 t / \gamma_1)}{y^2} \frac{(\sin y - y \cos y) \sin(r y / R_{NP})}{[(c \sin y - y \cos y)^2 + b^2 y^2 \sin^2 y]} dy \right\} \quad (18)$ |
| | $T_{Au}(r = 0, t)\vert _{t < 2\mu} = T_{ref} + \frac{R_{NP}^2 C_{abs} F}{k_{Au} V_{NP} 2\mu} \left\{ \frac{1}{3} \frac{k_{Au}}{k_m} + \frac{1}{6} - \frac{2b}{\pi} \int_0^\infty \frac{\exp(-y^2 t / \gamma_1)}{y^2} \frac{(\sin y - y \cos y)}{[(c \sin y - y \cos y)^2 + b^2 y^2 \sin^2 y]} dy \right\} \quad (19)$ |
| | $T_{water}(r, t)\vert _{t < 2\mu} = T_{ref} + \frac{R_{NP}^3 C_{abs} F}{r k_{Au} V_{NP} 2\mu} \left\{ \frac{1}{3} \frac{k_{Au}}{k_m} - \frac{2}{\pi} \int_0^\infty \frac{\exp(-y^2 t / \gamma_1)}{y^3} \frac{(\sin y - y \cos y) [b y \sin y \cos \sigma y - (c \sin y - y \cos y) \sin \sigma y]}{[(c \sin y - y \cos y)^2 + b^2 y^2 \sin^2 y]} dy \right\} \quad (20)$ |
| Cooling process | $T_{Au}(r, t)\vert _{t \geq 2\mu} = T_{ref} + \frac{2R_{NP}^3 b C_{abs} F}{r\pi k_{Au} V_{NP} 2\mu} \int_0^\infty \frac{\exp[-y^2(t - 2\mu)/\gamma_1] - \exp(-y^2 t / \gamma_1)}{y^2} \frac{(\sin y - y \cos y) \sin(r y / R_{NP})}{[(c \sin y - y \cos y)^2 + b^2 y^2 \sin^2 y]} dy \quad (21)$ |
| | $T_{Au}(r = 0, t)\vert _{t \geq 2\mu} = T_{ref} + \frac{2R_{NP}^2 b C_{abs} F}{\pi k_{Au} V_{NP} 2\mu} \int_0^\infty \frac{\exp[-y^2(t - 2\mu)/\gamma_1] - \exp(-y^2 t / \gamma_1)}{y^2} \frac{(\sin y - y \cos y)}{[(c \sin y - y \cos y)^2 + b^2 y^2 \sin^2 y]} dy \quad (22)$ |
| | $T_m(r, t)\vert _{t \geq 2\mu} = T_{ref} + \frac{R_{NP}^3 C_{abs} F}{r k_{Au} V_{NP} 2\mu} \left\{ \frac{1}{3} \frac{k_{Au}}{k_m} - \frac{2}{\pi} \int_0^\infty \frac{\exp[-y^2(t - 2\mu)/\gamma_1] - \exp(-y^2 t / \gamma_1)}{y^3} \frac{(\sin y - y \cos y) [b y \sin y \cos \sigma y - (c \sin y - y \cos y) \sin \sigma y]}{[(c \sin y - y \cos y)^2 + b^2 y^2 \sin^2 y]} dy \right\} \quad (23)$ |
| | $b = \frac{k_m}{k_{Au}} \sqrt{\frac{\alpha_{Au}}{\alpha_m}}, c = 1 - \frac{k_m}{k_{Au}}, \gamma_1 = \frac{R_{NP}^2}{\alpha_{Au}}, \sigma = \left( \frac{r}{R_{NP}} - 1 \right) \sqrt{\frac{\alpha_{Au}}{\alpha_m}} \quad (24)$ |

### Finite element simulation

All finite element simulations are conducted in COMSOL 5.3 software. The governing equations, boundary and initial conditions are described in previous section. The thermal interfacial resistance is described as the following equation:

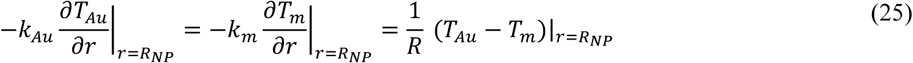

The volumetric heat generation by Gaussian pulse is defined as:

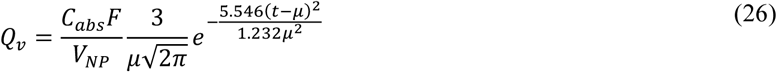

The medium domain is 10 times larger than the nanoparticle. The nanoparticle is located at the center of the medium domain. For gold nanorods (AuNRs), there are two important dimensions to check: axial dimension (z) and radial dimension (r). The origins of z and r are at the center of the nanorod.

The model is built in COMSOL 5.3 with 2D axisymmetric to decrease the computational time. The number of elements of the domain is in the range of 13461∼21517 depending on the shape of the nanoparticle. The maximum size is determined by mesh independence test and was set to be 1.5 nm. The time step is 0.1 ns and the total simulation time is 50 ns.

### Absorption cross section (C_abs_) calculation for AuNRs

The absorption cross sections (C_abs_) of AuNRs are calculated by MNPBEM tool box in MATLAB [33]. Because the C_abs_ of AuNRs is highly dependent incident light orientation, we calculate the average C_abs_ with varying the propagation and the polarization direction of incident light. The refractive index of water is constant 1.33 and dielectric function of gold is obtained from Johnson et al [34].

### Protein inactivation calculated by Arrhenius model

We assume protein inactivation is a two-state, first-order kinetic model with native (N) and inactivated (I) state:

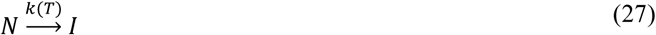

temperature-dependence of reaction rate can be described by Arrhenius model:

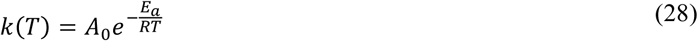

the inactivation of protein can be estimated by the following equation:

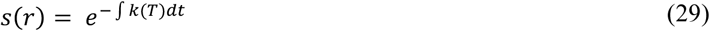

In this study, we first selected α-chymotrypsin as the model protein since our previous experiments demonstrated the feasibility of α-chymotrypsin photoinactivation by MH [25]. The pre-exponential factor (A_0_) of protein inactivation is 9.75×10^38^ s^-1^ and activation energy is 244.05 kJ mol^-1^ for α-chymotrypsin [19]. For other proteins, the Arrhenius kinetic parameters were derived from literature [35-41].

### Temperature-dependence of material properties

The material properties used in numerical and analytical solution are shown in table 3.

**Table 2.**
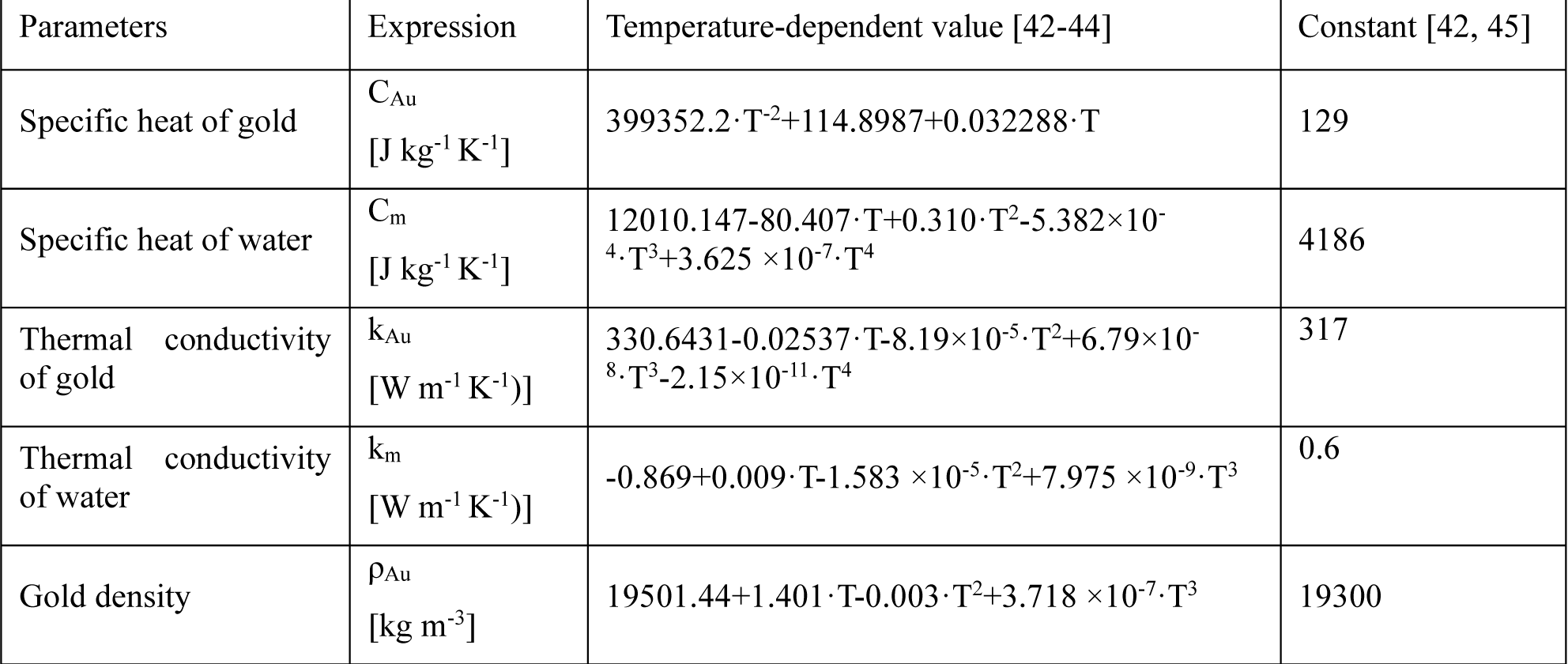
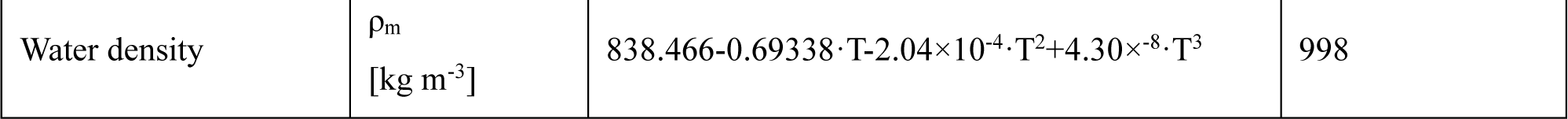
Temperature-dependence of material properties.

| Parameters | Expression | Temperature-dependent value [42-44] | Constant [42, 45] |
| --- | --- | --- | --- |
| Specific heat of gold | $C_{Au}$<br>[J kg<sup>-1</sup> K<sup>-1</sup>] | $399352.2 \cdot T^{-2} + 114.8987 + 0.032288 \cdot T$ | 129 |
| Specific heat of water | $C_m$<br>[J kg<sup>-1</sup> K<sup>-1</sup>] | $12010.147 - 80.407 \cdot T + 0.310 \cdot T^2 - 5.382 \times 10^{-4} \cdot T^3 + 3.625 \times 10^{-7} \cdot T^4$ | 4186 |
| Thermal conductivity of gold | $k_{Au}$<br>[W m<sup>-1</sup> K<sup>-1</sup>] | $330.6431 - 0.02537 \cdot T - 8.19 \times 10^{-5} \cdot T^2 + 6.79 \times 10^{-8} \cdot T^3 - 2.15 \times 10^{-11} \cdot T^4$ | 317 |
| Thermal conductivity of water | $k_m$<br>[W m<sup>-1</sup> K<sup>-1</sup>] | $-0.869 + 0.009 \cdot T - 1.583 \times 10^{-5} \cdot T^2 + 7.975 \times 10^{-9} \cdot T^3$ | 0.6 |
| Gold density | $\rho_{Au}$<br>[kg m<sup>-3</sup>] | $19501.44 + 1.401 \cdot T - 0.003 \cdot T^2 + 3.718 \times 10^{-7} \cdot T^3$ | 19300 |

**Table 2. continued**
|  |  |  |  |
| --- | --- | --- | --- |
| Water density | $\rho_m$<br>[kg m<sup>-3</sup>] | $838.466-0.69338 \cdot T-2.04 \times 10^{-4} \cdot T^2+4.30 \times 10^{-8} \cdot T^3$ | 998 |

**Table 3.**
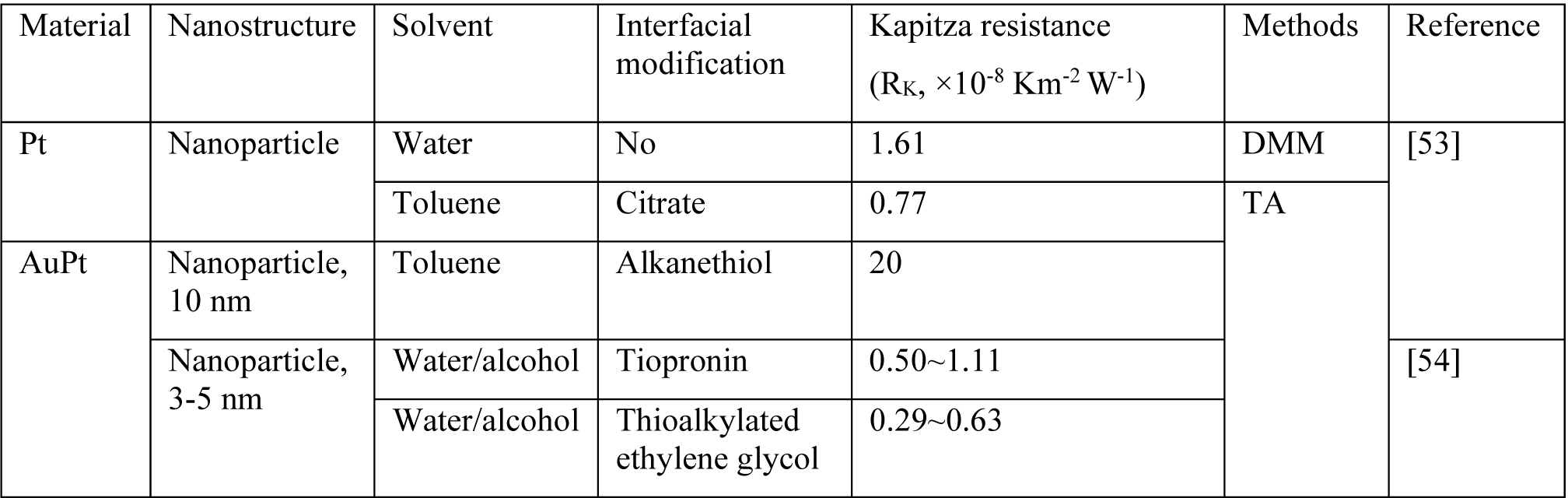
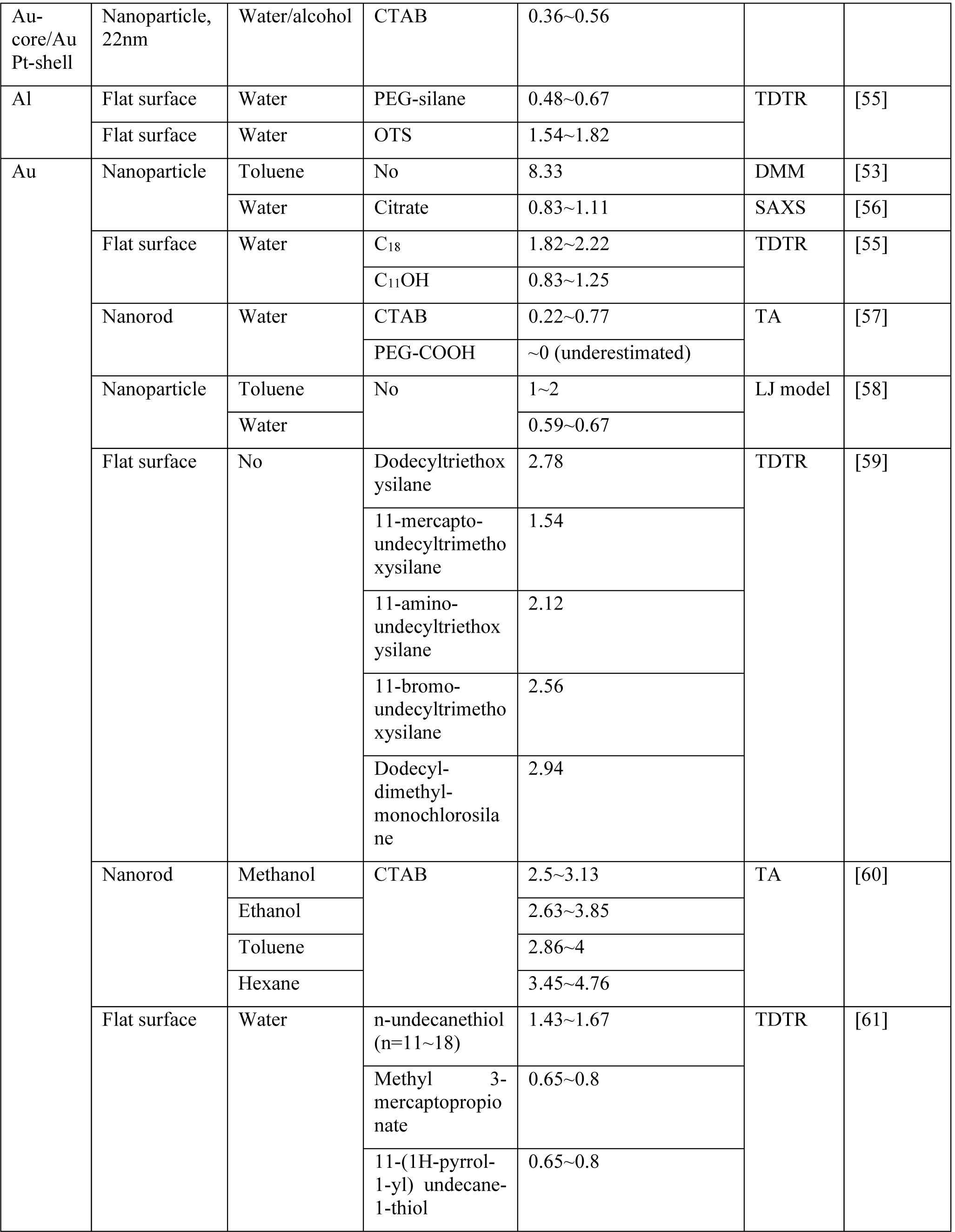
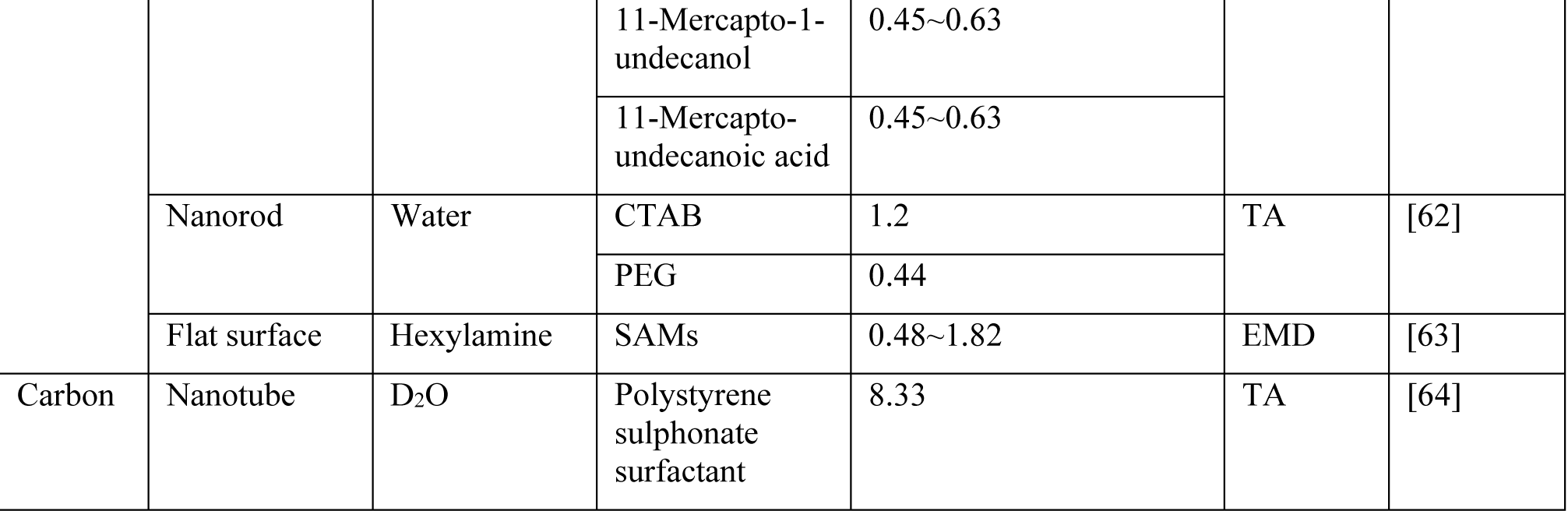
Summary of Kapitza thermal resistance of nanostructures from previous reports.

## Results

### 1. Localized heating *versus* collective heating

First, we developed a dimensionless parameter to differentiate localized heating versus collective heating, for molecular hyperthermia (MH) and hyperthermia, respectively. The effective thermal diffusion length in medium can be written as [46]:

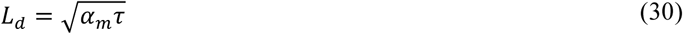

where *α*_*m*_ is medium thermal diffusivity, *τ* is heating duration. In a colloidal solution system, the mean inter-particle distance is given by following equation [47]:

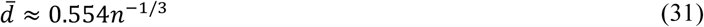

where n is the particle density (m^-3^). The ratio of L_d_ and 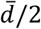 gives a dimensionless parameter *ξ*:

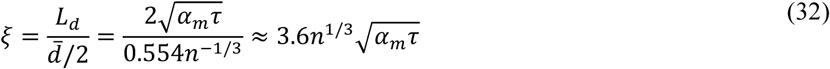

The dimensionless parameter *ξ* characterizes whether the heat from individual particles overlaps. When *ξ* is much smaller than 1, *i*.*e. ξ*<0.1, the thermal diffusion length is much smaller than inter-particle distance. Hence, the heat is localized around individual particles without overlapping, namely localized heating (Fig. 2A). The localized heating occurs when energy input duration is short and particle density is low. When *ξ* is close to 1, the heat diffusion length is close to inter-particle distance so that the heat from neighboring particles starts to overlap. With a continuous energy input, the heat overlap between particles will further develop and raises the temperature in a collective manner. This multiple particle heating can cause global tissue heating when *ξ* is much larger than 1 (*i*.*e. ξ*>10), known as collective heating [48]. With nanoparticle density of 10^15^ NP m^-3^, hyperthermia requires a relatively long laser irradiation (longer than 0.1s) to generate collective heating between particles and increases temperature of the whole tissue [49]. On the other hand, MH uses the nanosecond laser pulse to generate localized heating around particles and inactivates proteins, where *ξ < 0*.*01* (Fig. 2B). Therefore, the protein inactivation by MH falls well within a localized heating scenario.

**Figure 1.**
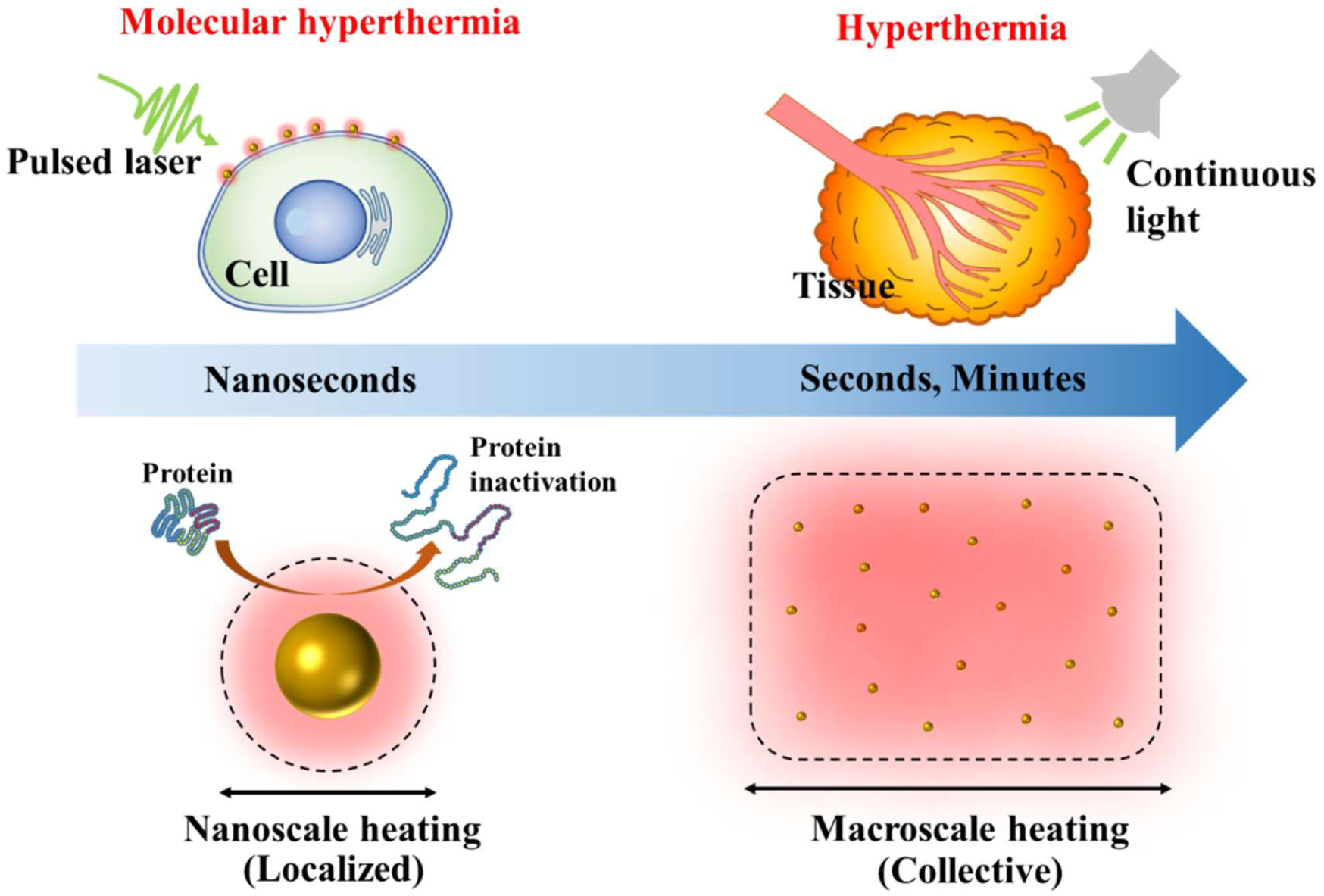
Comparison between molecular hyperthermia and hyperthermia. Molecular hyperthermia utilizes a nanosecond laser pulse to generate nanoscale heating (localized heating) and inactivate proteins without killing cells. On the other hand, hyperthermia uses macroscale heating (collective heating) of plasmonic nanoparticles induced by continuous irradiation (seconds to minutes) to kill tumor tissue.

**Figure 2.**
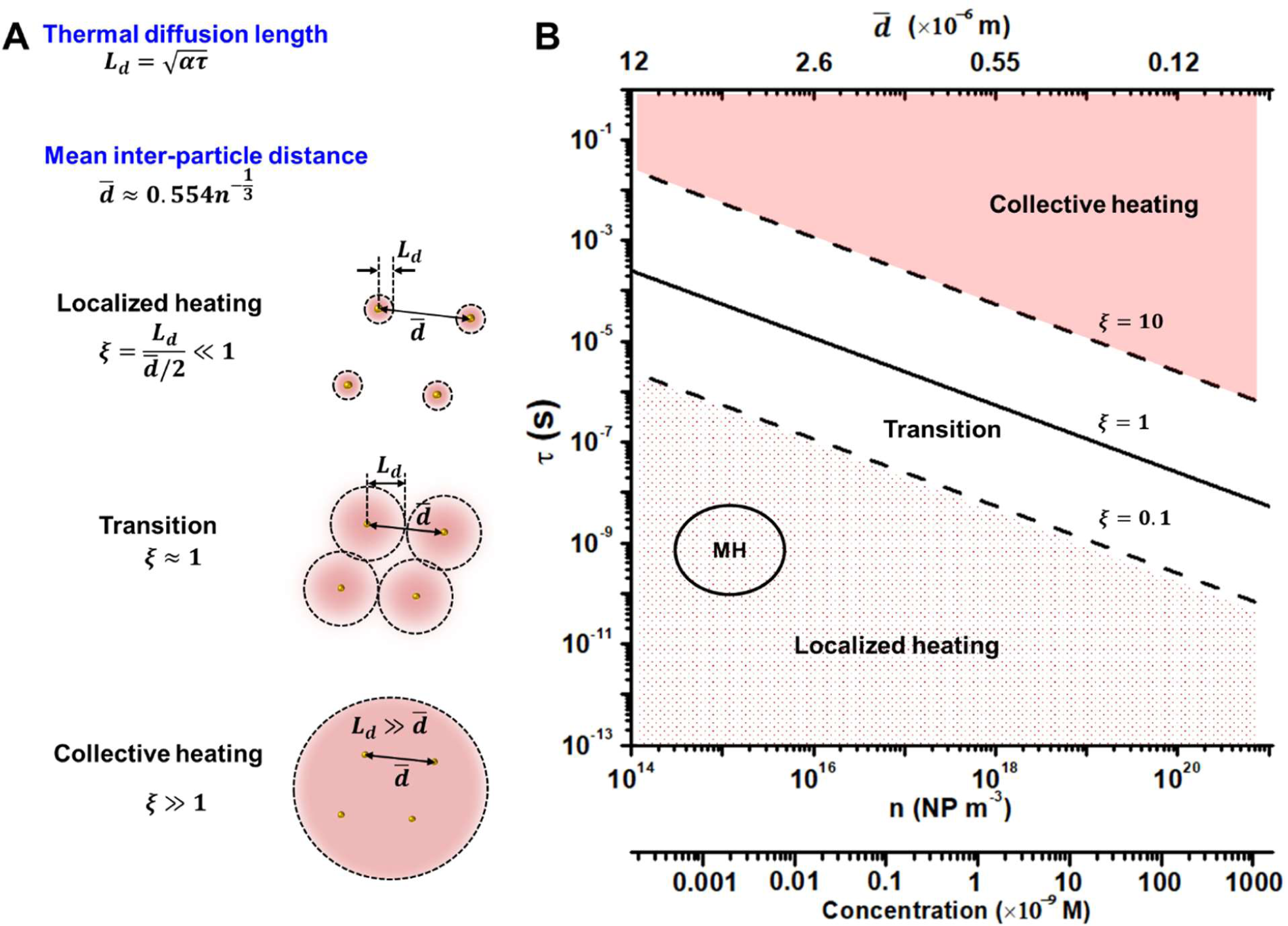
Dimensionless parameter to define conditions for localized and collective heating. (A) Schematic of localized heating versus collective heating. (B) Dimensionless parameter *ξ* to differentiate localized heating and collective heating. *τ* is the energy input duration and n is the nanoparticle density. The solid line indicates *ξ* =1, while the dashed lines indicate *ξ* =10 and 0.1, respectively. MH: molecular hyperthermia.

### 2. Numerical model validation

Second, we confirm the accuracy of the numerical model by comparing the numerical solution with the analytical solution. Fig. 3A-B illustrates the geometry and laser profile for the model. An AuNS with diameter of 30 nm (D_NP_ = 30 nm) is in the center of a spherical medium domain. The size of medium domain is 300 nm (D_medium_ = 300, Fig. 3A). The temperature of the medium domain boundary is set to 300 K (first-type boundary condition). For analytical solution, the AuNS is immersed in an infinite domain of water, which initial temperature is 300K. The single rectangular laser pulse (10 ns, laser intensity is 25.4×10^9^ W m^-2^) is used to heat the AuNS (Fig. 3B). Temperature profiles at three different positions from numerical simulation are compared with analytical solution (Fig. 3C). For both cases, the temperature rises quickly and returns to initial temperature (300 K) after 40 ns. The results demonstrate that the simulation time (50 ns) is sufficient for single particle to cool down. We also compared the numerical and analytical solution of temperature distribution at three different time points (Fig. 3D). Because the thermal conductivity of gold is significantly larger than water, the temperature distribution in AuNS is nearly uniform. The temperature difference between numerical solution and analytical solution is negligible. This suggests that the medium domain is large enough, and the first-type boundary condition (constant temperature) is satisfactory for this numerical simulation. The MH induced protein inactivation is calculated by the Arrhenius model. We assume that the protein distribution is homogeneous around the AuNSs. The protein inactivation profile was calculated along the radial direction after single pulse irradiation at 50 ns (Fig. 3E). The inactivation of proteins is localized around the AuNS and the border of inactivation zone is quite sharp. We define the impact zone as the distance from particle surface to where 50% of protein stay intact (Fig. 3E). Both numerical and analytical solutions give the same impact zone (∼10.5 nm, Fig. 3F). These results confirm the accuracy of the numerical method.

**Figure 3.**
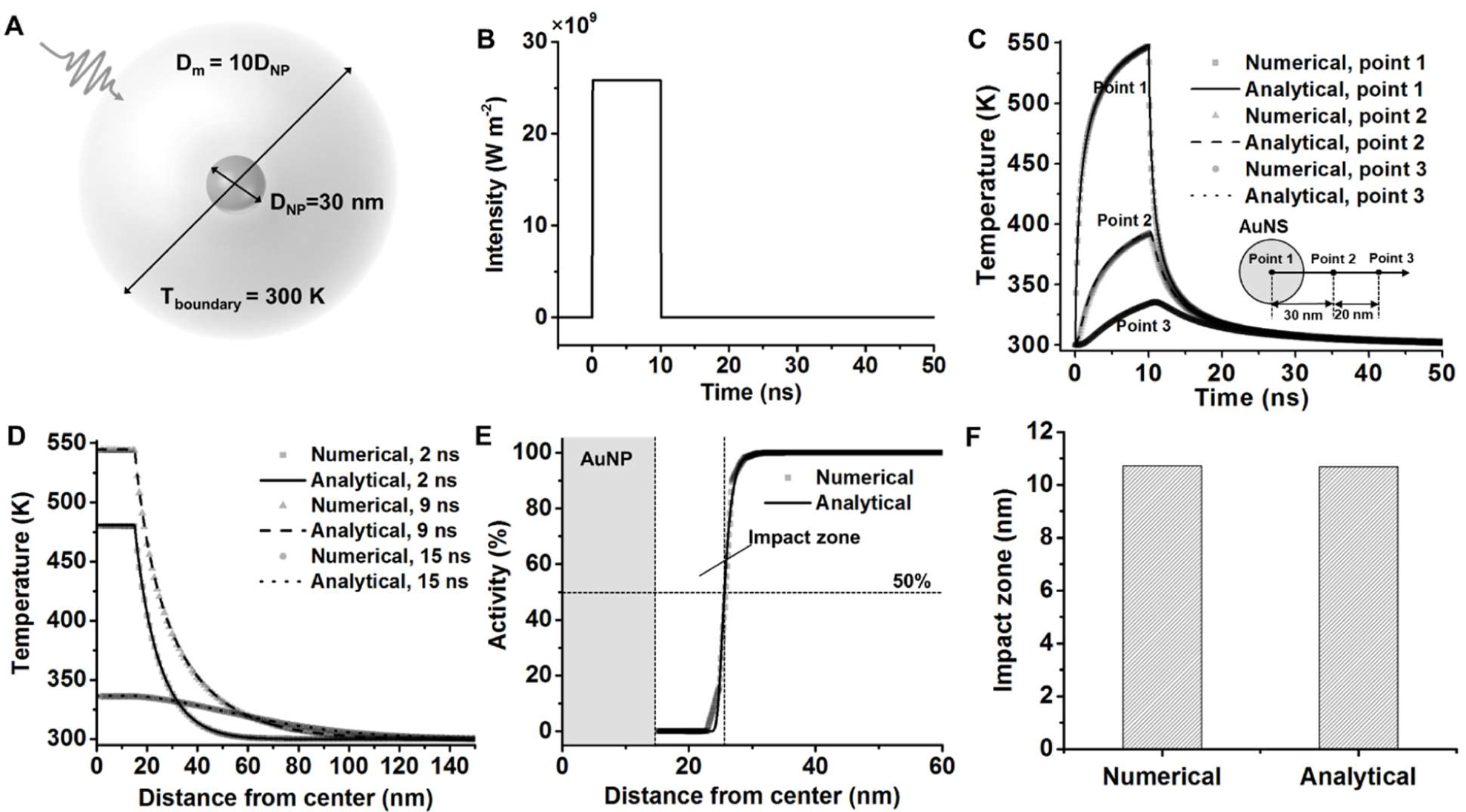
Numerical model validation. A) The model geometry for numerical simulation. D_NP_ is diameter of gold nanosphere (AuNS). D_m_ is diameter of medium domain and is set to be 10 times D_NP_. B) Rectangular laser intensity profile in both numerical and analytical solutions. C) Temperature profile of three different positions (illustrated by insert picture) by numerical and analytical solutions. D) Temperature distribution of three different time points by numerical and analytical solutions. E) Protein inactivation profile in the radial direction after laser irradiation. F) Impact zone predicted by the numerical method and analytical method.

### 3. Effects of pulse shape and duration

Secondly, we investigate the effect of pulse shape and pulse duration on MH. Here, we compare two pulse shapes, *i*.*e*. rectangular and Gaussian (bell). Also, we compare different durations of the Gaussian pulses. All laser pulses share the same fluence (254 J m^-2^, illustrated by shaded areas, Fig. 4A). All Gaussian pulses start at 0 ns time point, and their peak position varies from 5 ns to 20 ns depending on the pulse durations. Fig. 4B shows that the temperature response of AuNS varies according to pulse shapes and durations. The peak temperature of AuNS decreases when the pulse duration is stretched. For the rectangular shape pulse, the peak temperature of AuNS appears at the end of the pulse (10 ns). The protein activity distribution is then calculated based on the temperature profile (Fig. 4C). Although all the pulse energy inputs are same, pulses with longer duration inactivate less proteins than pulses with shorter duration. This is due to the nonlinear temperature-dependence of protein inactivation in the Arrhenius model. Pulses with shorter duration generates greater temperature rise and accelerates protein inactivation. Interestingly, the impact zone of rectangular pulse is same as the Gaussian pulse which peak position is at 10 ns (μ = 10 ns, Fig. 4D).

**Figure 4.**
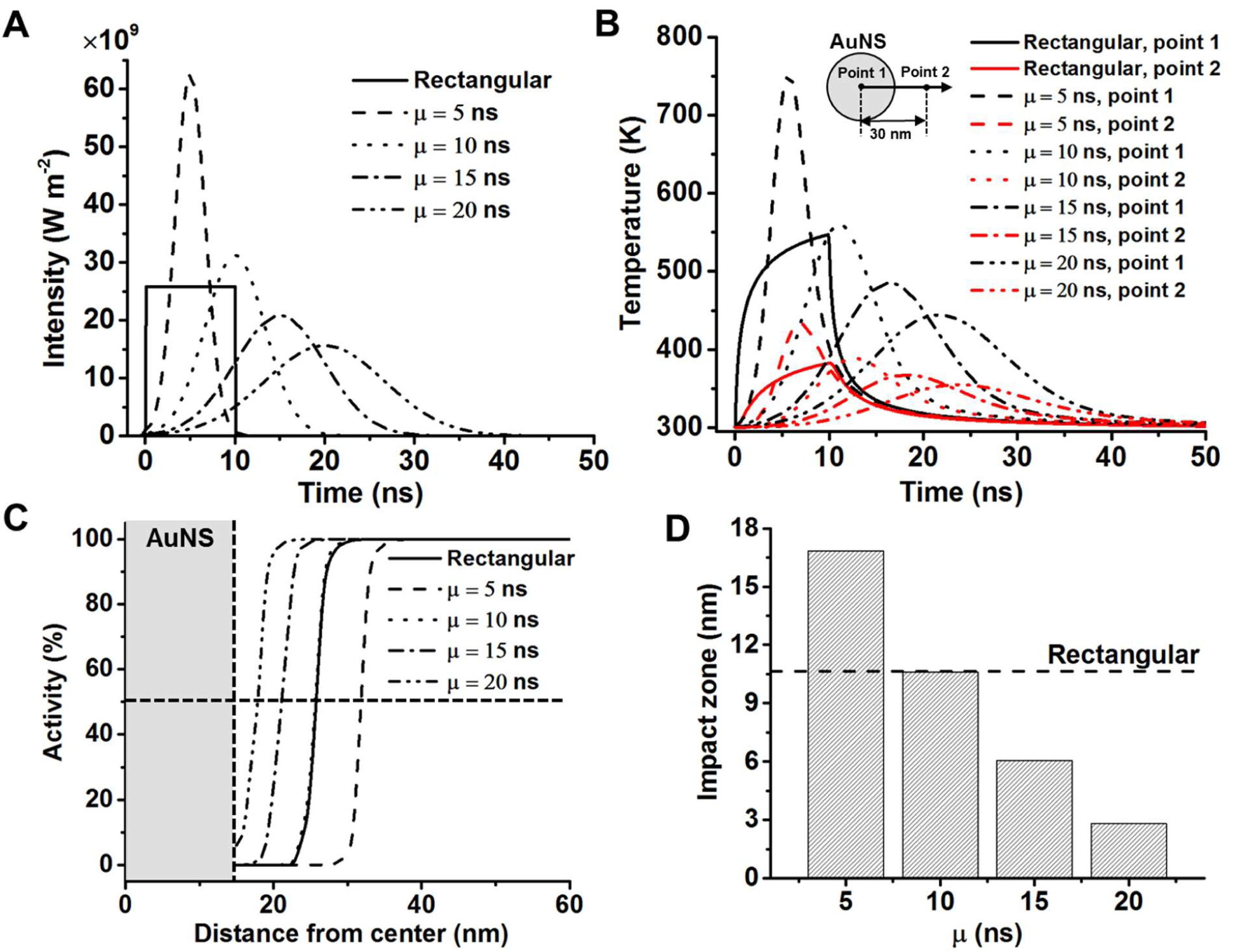
Effect of laser pulse shape and duration on molecular hyperthermia. A) The laser profiles with different shapes (Gaussian and rectangular) and different pulse durations. µ is the center of Gaussian pulse. All laser pulses share the same laser fluence (254 J m^-2^). B) Temperature profile of the gold nanosphere (AuNS) for different pulses. C) Protein activity distribution after different laser pulse. D) Effect of pulse duration on the impact zone. The dash line indicates impact zone of 10 ns rectangular pulse.

### 4. Effect of temperature-dependent material properties

Thirdly, we study the effect of temperature-dependent material properties (*k, C*_*p*_ and *ρ* in medium and AuNS) on the localized heating and protein inactivation. Fig. 5A shows that no obvious difference was observed for temperature-dependent density and specific heat. However, the temperature-dependent thermal conductivity significantly decreases the temperature profiles in both gold and medium. Consequently, the impact zone decreases 1.4 nm, 13% lower than the case with constant properties (Fig. 5B). If we consider the temperature-dependences for all material properties (labeled as [k, *ρ, C*_*p*_] (T)), the result is the same as the temperature-dependent thermal conductivity (labeled as k(T)). These results demonstrate that the change of thermal conductivity has a significant effect on MH. When applying MH on real biological environment such as cytoplasm, due to the high protein content in cells, the thermal conductivity of cytoplasm differs from pure water. The thermal conductivity of single live cells is measured to be 0.56∼0.58 Wm^-1^K^- 1^[50]. The protein concentration in the cytoplasm is about 100 mg mL^-1^ and this gives an estimated thermal conductivity of 0.57 W m^-1^ K^-1^.[51, 52] The impact zone increases slightly using cytoplasm thermal conductivity (11.3 nm) compared with using water properties (10.6 nm). Therefore, the MH can work more efficiently in cytoplasm than in pure water.

**Figure 5.**
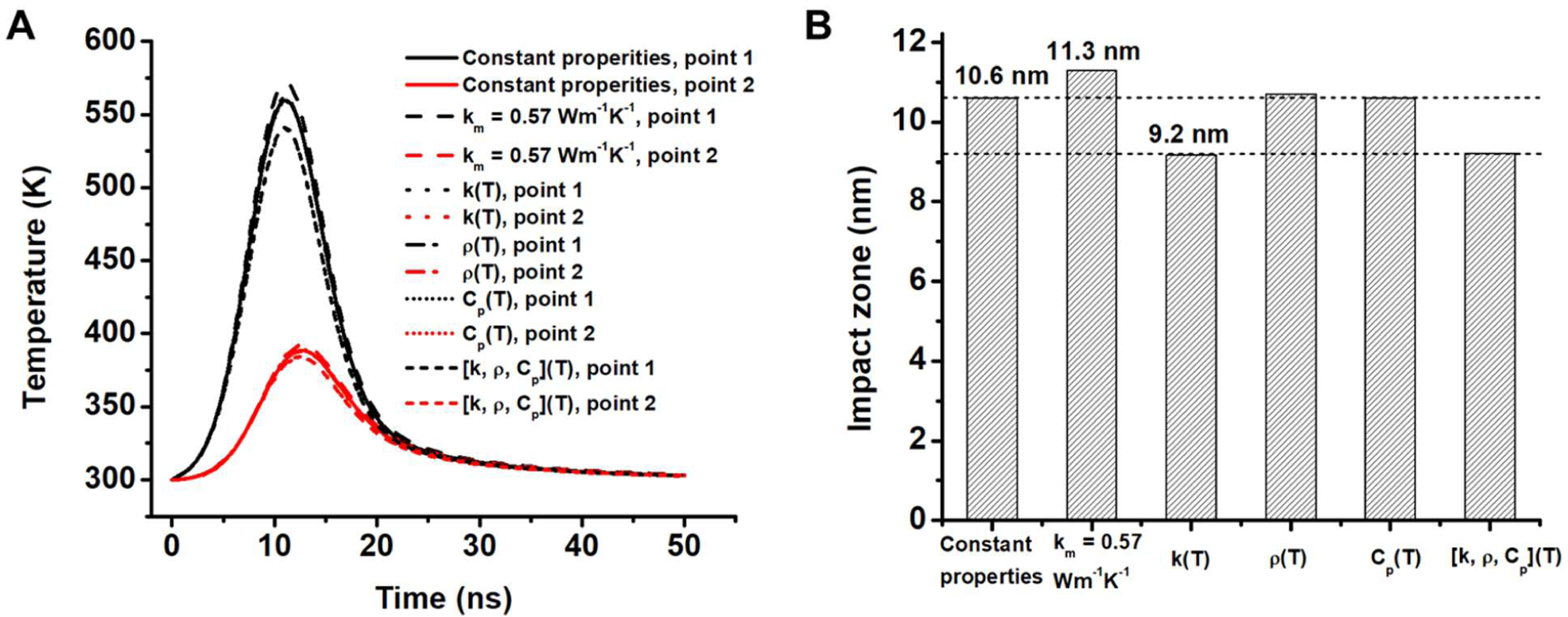
Effect of temperature-dependent material properties. (A) Comparison of temperature profile in different points for constant and temperature-dependent material properties. Point 1 indicates the center of the nanoparticle. Point 2 indicates the location in medium that 15 nm from the surface of the particle. (B) Comparison of impact zones for constant and temperature-dependent material properties and the cytoplasm. The “constant properties” indicates results with constant water properties. Specifically, for cytoplasm as medium, the thermal conductivity of medium (k_m_) is 0.57 Wm^-1^K^-1^, while other material properties are same as constant water properties. k (T), *ρ*(T) and C_p_(T) indicate that thermal conductivity, density and specific heating of materials are temperature dependent, respectively. The last group indicates results with temperature-dependences for all material properties ([k, *ρ, C*_*p*_] (T)).

### 5. Effect of thermal interfacial resistance

Next, we study the effect of thermal interfacial resistance (or Kapitza resistance, R_K_) at gold-medium interface on MH. R_K_ originates from the mismatch of phonon transport at the gold-medium interface, which can be significantly tuned by surface chemical modifications (Fig. 6A). In MH, the surface of AuNS is modified with polyethylene glycol (PEG) and antibodies. Thus, it is important to estimate the effect of R_K_ on MH in a wide range. In this study, a large range of R_K_ (1×10^−10^ Km^2^ W^-1^ to 1×10^−7^ Km^2^ W^-1^) is investigated to cover data reported previously (Table 3). Fig. 6B shows that R_k_ increases the temperature in the AuNS significantly. In contrast, medium temperature does not change significantly with changing R_K_. The medium temperature has a slight drop only when R_K_ is larger than 1×10^−8^ Km^2^W^- 1^ (Fig. 6B). The small drop in medium temperature is due to the heat flux decrease through the gold-medium interface (Fig. 6C). Consequently, the impact zone is smaller when R_K_ exceeds 1×10^−8^ Km^2^ W^-1^(Fig. 6D). However, when comparing our results with the R_K_ values reported by previous studies (marked by grey shade), the thermal interfacial resistance has a negligible effect on MH.

**Figure 6.**
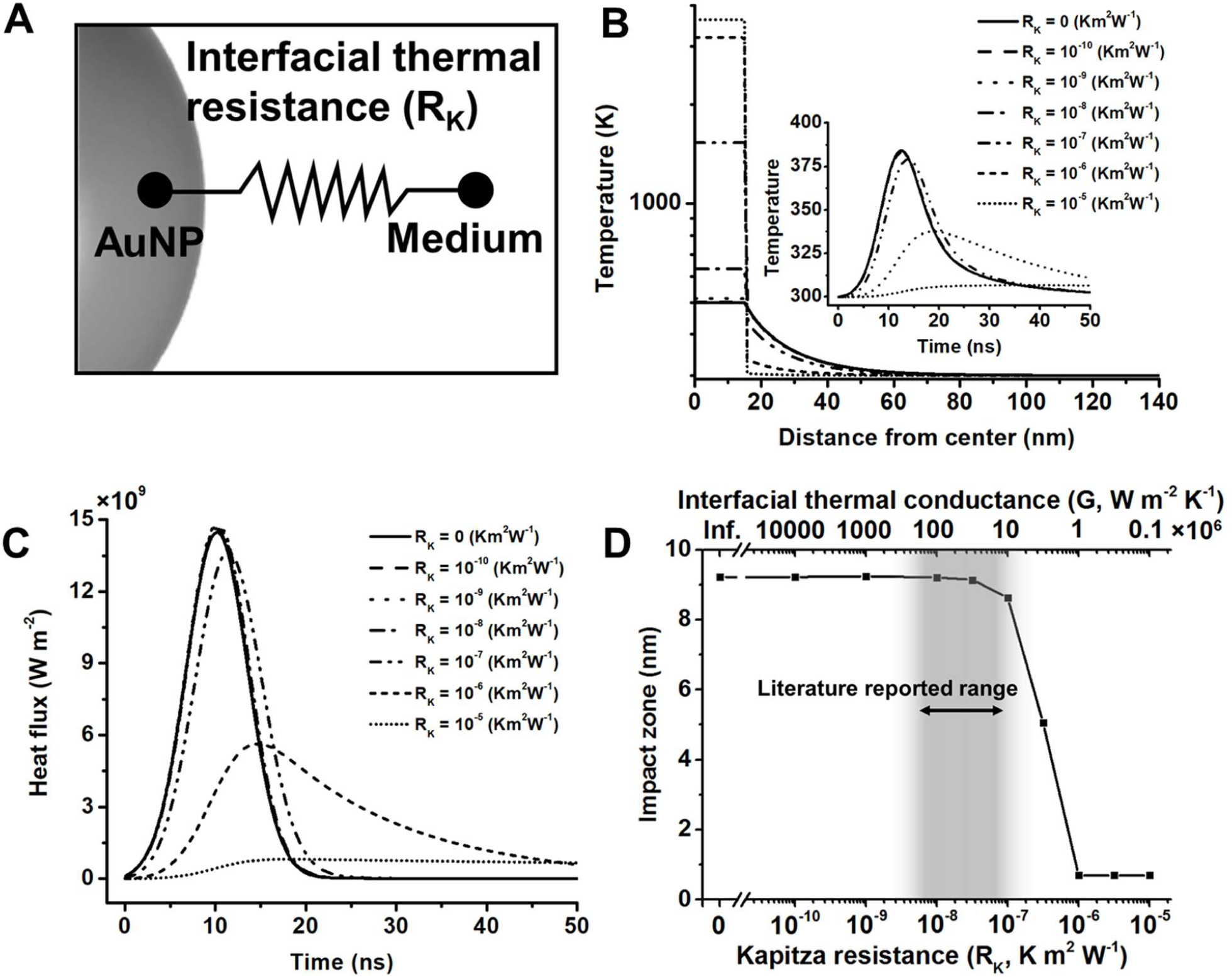
Effect of thermal interfacial resistance between gold and medium on protein inactivation. (A) Schematic of the interfacial resistance between AuNS and medium. (B) Temperature profile for different Kapitza resistances at t = 9 ns. Insert figures shows the temperature profile of medium at 30 nm from the nanoparticle center. (C) Heat flux at the gold-medium interface for different Kapitza resistance. (D) Impact zone for different Kapitza resistances. The interfacial thermal conductance (G) is the reciprocal of Kapitza resistance.

### 6. Effects of nanoparticle shape: nanosphere versus nanorods

Next, we investigate the effect of nanoparticle shape on MH. Gold nanorod (AuNR) is another type of nanoparticles that commonly used in biomedical related area. Biological tissue significantly absorbs and scatters visible light and is more transparent to near infrared light. AuNRs response to light in near-infrared range and offer better light penetration depth for biological tissue. To estimate the AuNR performance for MH, AuNRs with different aspect ratio are studied here (Fig. 7A). We keep two factors the same for all particles: particle volume (V_NP_) and volumetric heating rate (Q_v_). These conditions ensure the same amount of heat is generated among nanoparticles with different shapes. The results suggest that gold nanosphere (AuNS) has the highest temperature increase and nanoparticle temperature decreases with the aspect ratio (Fig. 7B&D-E). Compared with AuNS, AuNR has higher surface area and therefore enhances the heat dissipation to the surrounding medium, resulting in a lower nanoparticle temperature.

**Figure 7.**
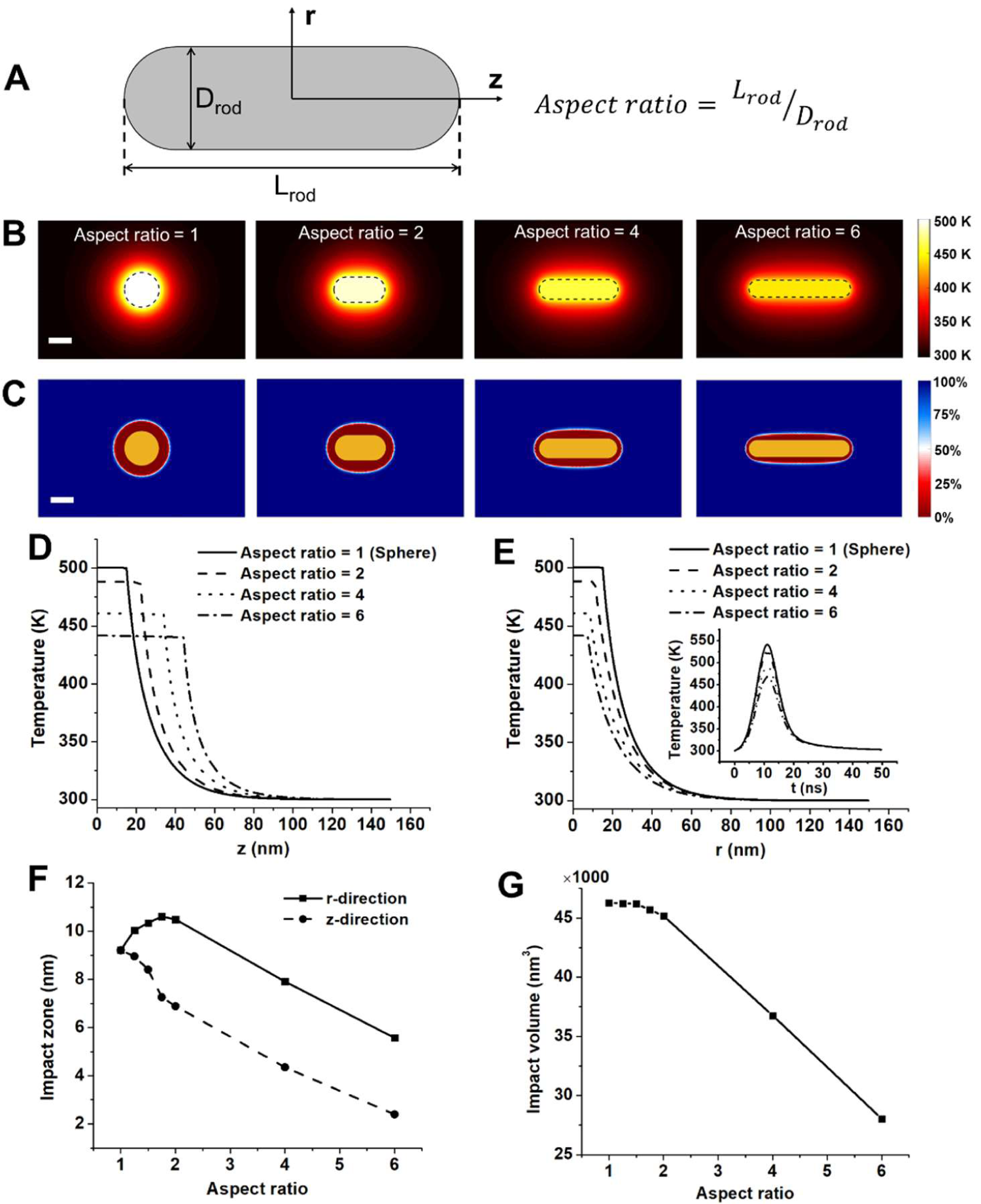
Effect of particle shape on molecular hyperthermia. (A) Geometry of gold nanorod (AuNR). Aspect ratio is defined as the ratio between length and radius of rod. All AuNRs in this study have the same volume. (B) Temperature distribution around AuNRs with different aspect ratios. (C) Protein activity distribution around AuNRs. 100% indicates all proteins are intact and 0% indicates all proteins are inactivated, the boundary of impact zone (50%) is shown in white color. (D) Temperature distribution in the axial direction for AuNRs with different aspect ratios. (E) Temperature distribution in the radial direction for AuNRs with different aspect ratios. Insert is gold temperature profile as a function of time. (F) Impact zone changes with aspect ratio in the axial and radial directions. (G) Impact volume for AuNRs with different aspect ratios.

We further investigated the protein inactivation for different shapes of gold nanoparticles (Fig. 7C&F-G). The impact zone in the axial direction (z-direction) decreases with higher aspect ratio. It is interesting that the impact zone in the radial direction (r-direction) firstly increases for nanorod with aspect ratio of 2 compared with spherical particle, but then decreases with aspect ratio from 2 to 6 (Fig. 7F). The impact zone in the r-direction reaches maximum value when aspect ratio is around 1.75. We further compared the impact volume for particle with different shapes, which is defined as the volume where proteins are inactivated by the localized heating. The impact volume decreases with increasing aspect ratio for AuNRs as a result of higher surface-to-volume ratio and heat dissipation rate than spherical nanoparticle (Fig. 7G). It is worth noting that we only considered constant volumetric heating generation in these comparisons. AuNRs have two plasmonic peak, corresponding to localized surface plasmon resonance (LSPR) from longitudinal and transverse modes. By adjusting the aspect ratio of AuNRs from 2 to 6, the longitudinal peak can be tuned from 600 nm to more than 1000 nm. Also, the longitudinal peak absorption is significantly higher than transverse peak (Fig. S1). Thus, by applying laser at longitudinal peak, AuNR would generate more heat than AuNS.

### 7. Effect of Arrhenius kinetic parameters on protein thermal denaturation

Lastly, we investigated different proteins using their respective kinetic parameters for thermal denaturation, *i*.*e*. the activation energy (E_a_) and the pre-exponential factor (A_0_). Here, we listed E_a_ and A_0_ of 19 different proteins in Table 4 from literature. Impact zones are calculated with the same condition in Fig. 5 (30 nm AuNS and temperature dependent material properties).

**Table 4.** Kinetic parameters and impact zones of different proteins. Kinetic parameters of the Arrhenius model are obtained from experimental measurements in the literature. The simulation is conducted with condition of 30 nm AuNS and temperature dependent material properties.

| Protein | $E_a$ (kJ mol<sup>-1</sup>) | $A_0$ (s<sup>-1</sup>) | Impact zone (nm) | Reference |
| --- | --- | --- | --- | --- |
| Amylase (malt) | 171.13 | $6.6 \times 10^{23}$ | <1 | [35] |
| Hemoglobin | 313.38 | $5.4 \times 10^{45}$ | 6.7 | [35] |
| Rennin | 371.54 | $6.4 \times 10^{57}$ | 13.2 | [35] |
| Egg albumin | 549.78 | $2.0 \times 10^{81}$ | 12.9 | [35] |
| Peroxidase (milk) | 772.37 | $1.6 \times 10^{114}$ | 15.6 | [35] |
| Hemolysin | 825.92 | $3.9 \times 10^{129}$ | 23.0 | [35] |
| Invertase (yeast) | 216.73 | $7.6 \times 10^{30}$ | 1.7 | [35] |
| | 459.40 | $5.9 \times 10^{52}$ | <1 | [35] |
| | 358.57 | $4.9 \times 10^{69}$ | >135 nm | [35] |
| | 308.36 | $4.6 \times 10^{45}$ | 7.4 | [35] |
| Opsin | 311.02 | $1.1 \times 10^{49}$ | 12.0 | [36] |
| Isorhodopsin | 386.18 | $6.9 \times 10^{57}$ | 11.0 | [36] |
| Rhodopsin | 415.59 | $1.7 \times 10^{62}$ | 11.3 | [36] |
| G actin | 231.67 | $2.8 \times 10^{33}$ | 2.9 | [37] |
| β-Trypsin | 275.55 | $7.9 \times 10^{41}$ | 7.7 | [38] |
| Horseradish peroxidase | 159.78 | $5.3 \times 10^{24}$ | 1.4 | [39] |
| Catalase | 254.78 | $2.1 \times 10^{38}$ | 6.1 | [40] |
| α-chymotrypsin | 244.31 | $9.8 \times 10^{38}$ | 9.3 | [19] |
| P-glycoprotein | 161.25 | $1.8 \times 10^{23}$ | <1 | [41] |

Most of proteins in our study can be inactivated by MH with an impact zone from 1-20 nm. Some proteins in the list are interesting in the context of biomedical engineering and sciences. For example, opsin is a G protein coupled receptor (GPCR) and plays an important role in sensory neurons for vision. GPCRs is a large protein family and contains many potential drug targeting sites for disease treatments. In our previous study, we have investigated protease activated receptor 2 (PAR2), which is also a GPCR [26]. Although kinetic parameters vary from protein to protein, impact zones of opsin and related receptors suggest MH can be a useful method to inactivate GPCRs. On the other hand, the activity of some proteins could be insensitive to MH. For example, P-glycoprotein (P-gp) can be overexpressed by cancer cells and is related to multidrug resistance in cancer chemotherapies. Many researches tried to inhibit activity of P-gp in cancer cells as a method to over the tumor multidrug resistance. However, we observed that the impact zone of P-gp is below 1 nm, meaning that they cannot be inactivated under current condition of MH. This may be due to that higher temperature is required, or the measured Arrhenius parameters need further validation.

Since different proteins are characterized by different Arrhenius parameters, we then systemically investigated a range of Arrhenius parameters on MH (Fig. 8). There are two interesting findings. Firstly, localized protein inactivation only occurs when the protein’s kinetic parameters are in the triangle shaded area shown in Fig. 8A. We plotted all protein kinetic parameters from Table 4 in the figure and most are localized in the “triangle area”. Previous studies reported correlation between E_a_ and A_0_ and are also displayed in Fig. 8A [65, 66]. Above the triangle area, the impact zone is very large, and all proteins get inactivated even at the boundary of the computational domain (which is held at constant temperature, 300 K). This suggests that the combination of E_a_ and A_0_ gives a thermally unstable situation. Below the triangle area, the impact zone is less than 1 nm. The combination of E_a_ and A_0_ does not lead to any protein inactivation. Secondly, we observed that the impact zone seems to correlate with the activation energy E_a_ (Fig. 8B). The protein with E_a_ smaller than 200 kJ mol^-1^ has a very small impact zone. The impact zone increases when E_a_ is larger than 200 kJ mol^-1^. The results also demonstrate that the protein inactivation by MH is highly localized around the nanoparticle as the impact zone hardly exceeds 20 nm.

**Figure 8.**
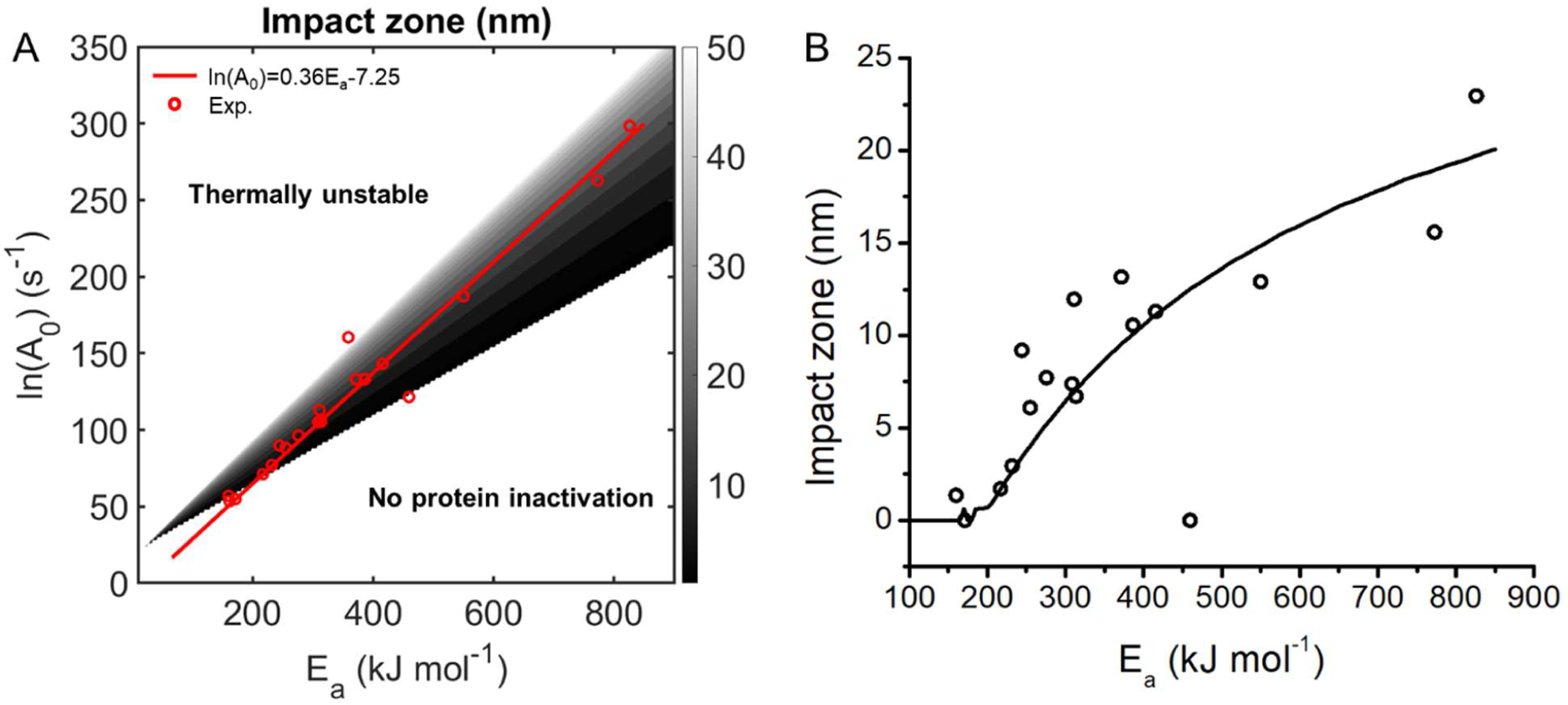
Impact zones for different proteins. (A) Contour plot of impact zone as a function of E_a_ and A_0_. The shaded region suggest that molecular hyperthermia can inactivate proteins effectively. The red dots are experimental measurements of E_a_ and A_0_ for different proteins from Table 4. The solid line indicates the empirical correlation between E_a_ and A_0_ [65]. (B) Impact zones calculated based on the E_a_-A_0_ correlation (black line) and real protein kinetic parameters (black dots). The thermally unstable proteins are not plotted on the figure since they have unrealistic impact zones.

## Discussion

In this study, we built a computational model to study the protein inactivation behavior during MH. We studied important factors that affect the protein inactivation in MH including laser pulse shape, duration, temperature-dependent material properties, thermal interfacial resistance, nanoparticle shape, and different proteins. The results provide a guideline in designing and applying MH for various biomedical applications. There are several interesting aspects that have emerged from our studies.

### Material properties in the nanoscale

The material properties utilized here are all obtained from bulk materials. However, the physical properties of material may vary in the nanoscale. For example, the melting temperature of AuNS drops significantly compared with bulk gold when the particle size is smaller than 10 nm [45]. Since we use 30 nm AuNS in our study, the melting point of particles in our study can be considered the same as bulk gold (1337K). Also, we limited our laser energy to ensure the nanoparticle temperature below gold melting point. Therefore, we did not consider the phase change effects in this study. However, AuNS could undergo phase change process at high laser fluence and possibly reshaping and fragmentation. Besides melting temperature, the thermal conductivity is also reported to be highly dependent on material size when it comes to nanoscale. Chen et al. estimated the thermal conductivity for 3 nm AuNS to be 24.14 W m^-1^K^-1^, an order magnitude lower than bulk gold.[67] In our previous study, we compared finite element model and molecular dynamics model to investigate the material properties of both gold and medium in nanoscale [25]. We found an obvious mismatch between two models when we use bulk material properties in FEM. This indicates that the size-dependence of material properties needs to be considered in computational methods. However, a lot remains unknown in this area and further experimental and numerical studies are required to elucidate the heat transfer process in the nanoscale.

### Protein inactivation in wide temperature range

In this study, we adopted a conventional Arrhenius model for protein inactivation process in which both pre-exponential factor (A_0_) and activation energy (E_a_) are independent of temperature. This hypothesis is valid for low temperature range. However in a wider temperature range, the activation energy and pre-exponential factor both vary with temperature as demonstrated by our previous study [27]. Also, the protein inactivation process is assumed to a first-order, irreversible process in this study. The thermal inactivation of protein is a much more complicated process and is highly dependent on the protein structure [68].

## Conclusions

In this study, we numerically investigate protein photoinactivation during MH by combining the heat transfer model with the chemical reaction model. Impact zone is defined to quantify the protein inactivation efficiency. Firstly, we studied the effect of pulse shape and duration and found that stretching the laser pulse duration reduces impact zone with the same laser energy. Secondly, we demonstrated that temperature-dependent material density and specific heat have negligible effect on MH, while temperature-dependent thermal conductivity decreases the impact zone and the low thermal conductivity of cytoplasm increases the impact zone. Thirdly, thermal interface resistance has a limited impact on MH below 1×10^−8^ Km^2^ W^-1^, a value larger than reported values for interface resistance. Fourthly, the nanosphere has larger impact volume than nanorods with the same heat generation. Lastly, the protein Arrhenius parameters have significant influence on the impact zone. In summary, this work provides an analytical interpretation of protein photoinactivation by MH and offers a clear guideline to design MH for biomedical applications.

## Acknowledgements

Research reported in this work was partially supported by National Institute of General Medical Sciences (NIGMS) of the National Institutes of Health under award number R35GM133653, 2019 Collaborative Sciences Award from American Heart Association under award number 19CSLOI34770004, and High-Impact/High-Risk Research Award from Cancer Prevention and Research Institute of Texas under award number RP180846. The content is solely the responsibility of the authors and does not necessarily represent the official views of the funding agencies.

## Nomenclature

AuNR = gold nanorod

AuNS = gold nanosphere

A_0_ = pre-exponential factor in Arrhenius model

C_abs_ = absorption cross section area, m^2^

Cht = α-chymotrypsin

C_p_ = specific heat, J kg^-1^ K^-1^

CTAB = cetyl trimethylammonium bromide

DMM = diffuse-mismatch model

D_NP_ = diameter of nanoparticle, 2R_NP_, m

D_rod_ = diameter of rod

D_m_ = diameter of medium domain, 2R_m_, m

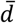= mean inter-particle distance, m

EMD = equilibrium molecular dynamics

E_0_ = activation energy in Arrhenius model

F = laser fluence, J m^-2^

G = interfacial thermal conductance, W K^-1^ m^-2^

GPCR = G protein coupled receptor

k = chemical reaction rate constant, s^-1^

k_Au_ = thermal conductivity of gold, W m^-1^ K^-1^

k_m_ = thermal conductivity of medium, W m^-1^ K^-1^

LJ = Lennard-Jones

L_d_ = thermal diffusion length. m

L_rod_ = length of rod

LSPR = localized surface plasmon resonance

MD = molecular dynamics

MH = molecular hyperthermia

OTS = Octadecyltrichlorosilane

PEG = polyethylene glycol

P-gp = P-glycoprotein

Q_v_ = volumetric heating rate, W m^-3^

R = the gas constant, 8.314 J K^-1^ mol^-1^

R_K_ = Kapitza resistance, Km^2^ W^-1^

SAM = self-assembly monolayer

SAXS = small angle x-ray scattering

s = survival fraction of protein

T = temperature, K

TA = transient absorption measurement

TDTR = time-domain thermoreflectance

t_0_ = laser duration, s

u(t) = unit step function

V_NP_ = nanoparticle volume, m^3^

α = thermal diffusivity, m^2^ s^-1^

µ = center position of the pulse, s

2µ = pulse duration, s

*ξ* = dimensionless parameter, 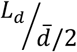

## Supporting information

**Figure S1.**
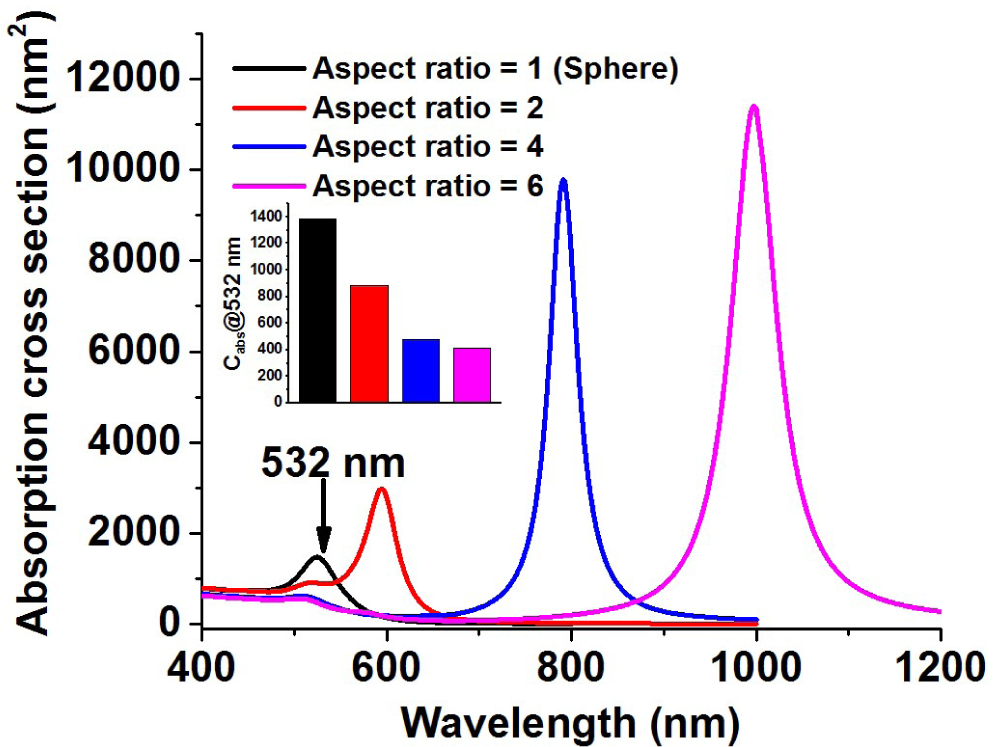
Calculated absorption cross section (C_abs_) for gold nanosphere and nanorods with different aspect ratio. Insert: comparison of C_abs_ at 532 nm laser wavelength.

**Figure S2.**
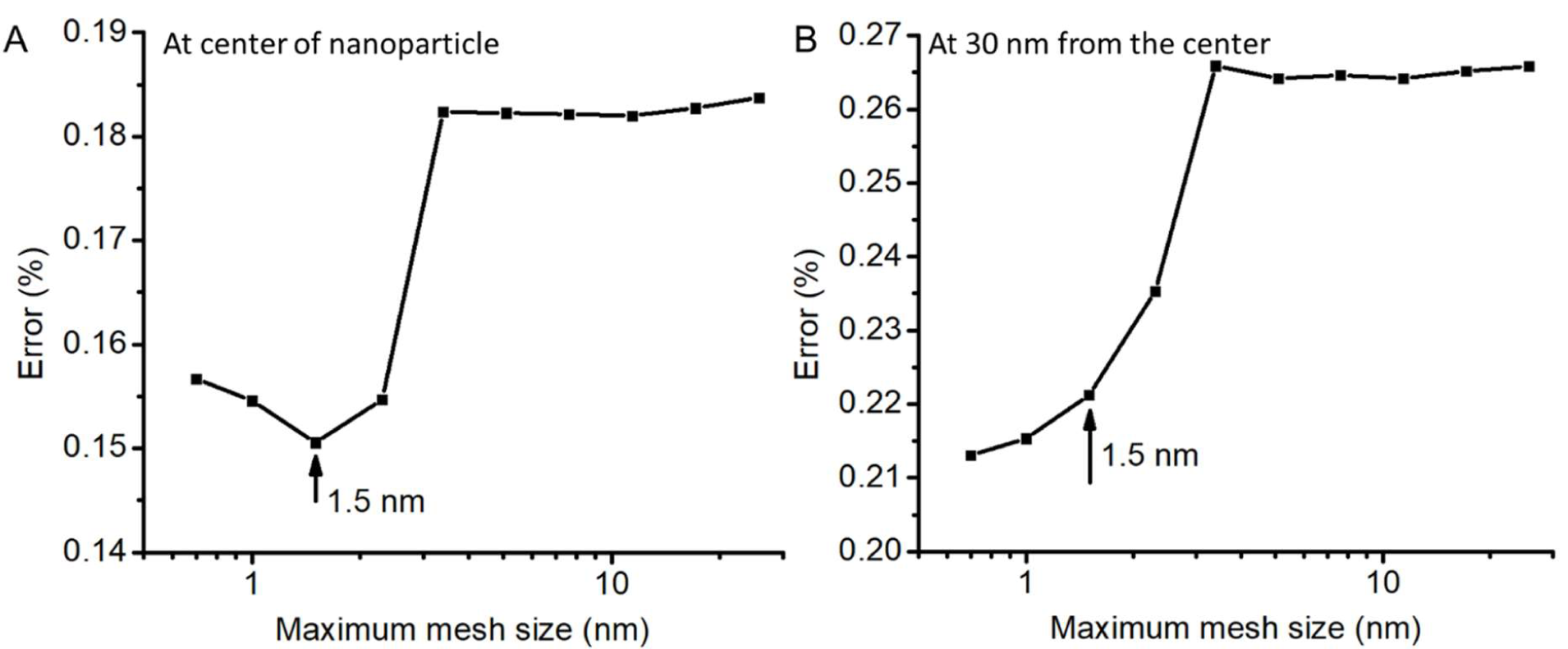
Mesh independence test. The temperatures at two different points were checked while varying the maximum mesh size. The error was calculated by comparing the numerical solution with analytical solution. Mesh size of 1.5 nm was used for this study.

